# Sensorimotor Confidence for Tracking Eye Movements

**DOI:** 10.1101/2023.04.28.538675

**Authors:** Alexander Goettker, Shannon M. Locke, Karl R. Gegenfurtner, Pascal Mamassian

**Affiliations:** Abteilung Allgemeine Psychologie, Justus-Liebig University Giessen, Giessen, Germany; Laboratoire des systèmes perceptifs, Département d’études cognitives, École normale supérieure, PSL University, CNRS, 75005 Paris, France

## Abstract

For successful interactions with the world, we often have to evaluate our own performance. Such metacognitive evaluations have been studied with perceptual decisions, but much less with respect to the evaluation of our own actions. While eye movements are one of the most frequent actions we perform, we are typically unaware of them. Here, we investigated if there is any evidence for metacognitive sensitivity for the accuracy of eye movements. Participants tracked a dot cloud as it followed an unpredictable sinusoidal trajectory for six seconds, and then reported if they thought their performance was better or worse than their average tracking performance. Our results show above chance identification of better tracking behavior across all trials and also for repeated attempts of the same target trajectories. While the sensitivity in discriminating performance between better and worse trials was stable across sessions, for their judgements participants relied more on performance in the final seconds of each trial. This behavior matched previous reports when judging the quality of hand movements, although overall metacognitive sensitivity for eye movements was significantly lower. Together, these results provide an additional piece of evidence for sensorimotor confidence, and open interesting questions about why it differs across actions and how it could be related to other instances of confidence.

**Statement of Relevance:** In everyday life, it is often critical to be able to evaluate the quality of our own cognitive decisions and actions. However, one of our most frequent actions often does not even reach our awareness: eye movements. We investigated whether observers were able to successfully judge the accuracy of their eye movements when tracking a cloud of dots that followed an unpredictable trajectory. We found that observers were able to distinguish good from bad trials, although sensitivity was lower compared to equivalent previous reports when judging the quality of hand movements. These results add an item to the growing list of our metacognitive abilities, but critically for eye movements that we are typically unaware of. They lead to important novel questions about why metacognitive abilities differ across decisions or different types of actions, and what the potential components of metacognitive abilities might be.

## Introduction

When was the last time you moved your eyes? The answer is probably only a few hundred milliseconds ago, when your eyes jumped from word to word while reading this text. Despite the importance of eye movements for visual perception (Findlay & Gilchrist, 2003; Gegenfurtner, 2016; Schütz et al., 2011), we usually only have limited awareness of when and how our eyes move (Goettker et al., 2018; Nieuwenhuis et al., 2001; van Zoest & Donk, 2010). Eye movements are important since they allow us to place the high-acuity foveal region on visual elements of interest. Tracking moving objects in the scene also allows for better estimates about the object’s properties (Lisberger, 2015; Schütz et al., 2009; Spering et al., 2011). Eye movements are conducted with high precision and low latency (Carpenter, 1988; Leigh & Zee, 2015; Liston & Stone, 2014). Yet, eye movements are not perfect, and will incur some amount of error due to different sources of noise (van Beers, 2007). Errors in eye movements can be registered by the brain as shown by oculomotor adaptation, where these error signals drive motor learning (for a review, see Pélisson et al., 2010). But, can error signals be used by the observer to judge the accuracy of their own eye movements?

Recent research indicates that humans can estimate their error to report sensorimotor confidence for visually-guided hand movements (Locke et al., 2020; Mole et al., 2018). Sensorimotor confidence is a subjective report of performance in a task that takes into account 1) the quality of the perceptual information, 2) the quality of the motor execution, and 3) the sensorimotor goal (Locke et al., 2020). In these studies, participants used their hands to control an input device in a computerized game. Afterwards, they made a subjective judgement about their performance in the task (i.e., a sensorimotor confidence report about their accuracy), which to some degree agreed with their true objective performance. But can we generalize these studies to eye movements? There are reasons to doubt that sensorimotor confidence for eye movements will be similar, since their function (Land & Hayhoe, 2001) and the neuroanatomy of oculomotor circuits (Leigh & Zee, 2015) substantially differ from that of voluntary hand movements.

The few studies that have investigated the subjective experience of eye movements have focused on motor awareness, not sensorimotor confidence. Motor awareness is the knowledge one has about whether their eye has moved. In contrast, sensorimotor confidence specifically considers the quality of the perceptual information and the goals of the movement. However, motor awareness can still be a necessary component for assessing the quality of motor execution for sensorimotor confidence: how could one assess the quality of our eye movements if we cannot monitor that our eyes moved? While we are usually not aware of how our eyes move, we can bring them under cognitive control and voluntarily direct and keep our gaze at a certain visual stimulus, for example a fixation cross (Gegenfurtner, 2016; Thaler et al., 2013), or look back at a remembered location (Pierrot-Deseilligny et al., 1991). Eye movement latencies can also be altered by reinforcement learning (Vullings & Madelain, 2018) and are under discriminative control depending on the visual consequences (Vullings & Madelain, 2019). Recently, Vencato and Madelain (2020) could even show that after training, observers were able to estimate their own saccade latency with an accuracy of about 40 ms. While these results clearly show that observers can sometimes control and monitor some information about their eye movements, other results indicate that eye movements sometimes escape voluntary control. In particular, there are circumstances where the eyes can react to things that are imperceivable (Spering & Carrasco, 2015). Tavassoli and Ringach (2010) demonstrated that the eyes react to fluctuations in target velocity, that are perceptually not detectable. If observers in this task were completely aware of their movements, they could have been able to use those as a cue to detect the velocity fluctuation. These results speak for limited motor awareness of eye movements and therefore possible limited sensorimotor confidence. However, it is important to remember that corrections without awareness of changes in the target position are also possible for hand movements (Goodale et al., 1986; Prablanc & Martin, 1992). Still, for hand movements, decisions seem to be based on knowledge about their own variability (Trommershäuser et al., 2008), and as reviewed above, reliable sensorimotor confidence has been reported (Locke et al., 2020; Mole et al., 2018).

Here we wanted to directly address the question whether observer can judge the accuracy of their own eye movements by measuring sensorimotor confidence for tracking movements. We used a paradigm similar to Locke et al. (2020), thereby also allowing us to directly compare the results of the present study with these previous results. Participants had to track the trajectory of an unpredictably moving target, whose position was only indicated by randomly sampled dots (see Figure 1). Subsequently, participants had to judge whether they tracked the target more accurately than their average performance. We preregistered the study and analysis (see Methods for more details). We expected that observers would show sensorimotor confidence for eye movements as reflected in an above chance performance to sort better from worse eye movements. In addition, we expected to find this confidence ability not only across all trajectories in the experiment, but also within repeats of the same trajectories. Furthermore, we were interested in whether sensorimotor confidence is stable within trials and across different sessions across days.

**Figure 1:**
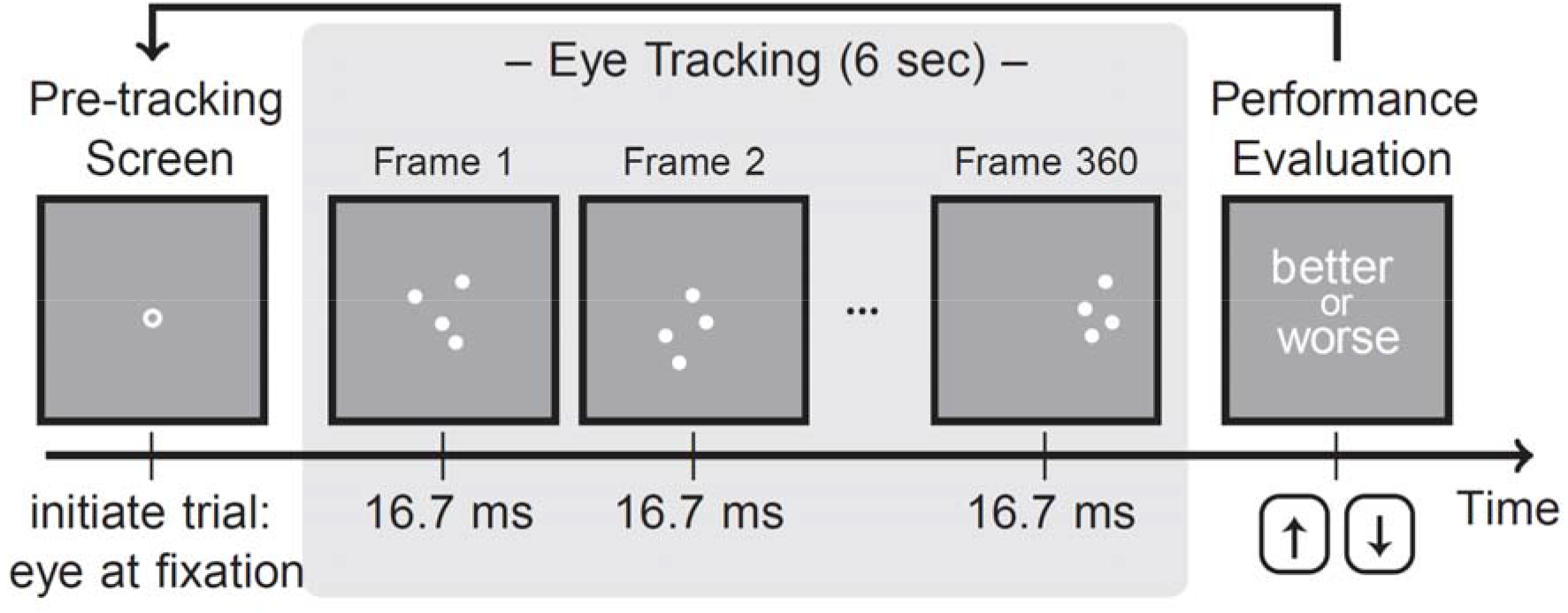
Trial Structure. Participants initiated the trial with a fixation check, then tracked the center of a moving dot cloud for 6 seconds. Four dots were drawn each frame from a 2D Gaussian distribution whose horizontal position followed an unpredictable sum-of-sinusoids trajectory. Sensorimotor confidence was reported as a binary better/worse performance evaluation, where participants reported their belief about their tracking accuracy relative to their personal average level of tracking accuracy.

## Methods

### Participants

Thirty participants (mean age of 24.56 years old, 21 female) with normal or corrected-to-normal vision took part in the study. All participants but one author were naive to the design of the experiment. Testing was conducted in accordance with the ethics requirements of the ethics committee of the Justus-Liebig University Giessen. Participants received details of the experimental procedures and gave written informed consent prior to the experiment and were paid 8 Euros per hour for taking part in the experiment.

### Setup

Stimuli were displayed on a Philips Brilliance 288P Ultra Clear monitor (60 × 32 cm, 3840 × 2160 pixel, 60 Hz). Participants sat 70 cm from the monitor with their head stabilized by a chin rest. Gaze was recorded from one eye with a desk-mounted eye tracker (EyeLink 1000 Plus, SR Research, Kanata, ON, Canada) at a sampling frequency of 1000 Hz. Before each block a nine-point calibration was used, and an additional drift check was performed at the start of each trial.

All confidence judgements were entered on a standard computer keyboard. The experiment was conducted using custom-written code in MATLAB version R2021b (The MathWorks, Natick, MA), using Psychtoolbox version 3.0.16 (Brainard, 1997; Kleiner et al., 2007; Pelli, 1997).

### Stimuli

We presented a horizontally-moving dot cloud to be gaze-tracked for 6 seconds (see Figure 1). On each frame, four static white dots (diameter: 0.25 deg) were sampled from an 2D circularly-symmetric Gaussian distribution (standard deviation: 2 deg; centered vertically) and presented on a mid grey background. Dots disappeared and were replaced every frame, giving the stimulus a twinkling quality. The tracking target was the mean of this invisible dot-generating distribution, which followed one of ten pre-computed horizontal trajectories. The same ten trajectories were used for all participants and presented multiple times but with new dots sampled for each repeat. The trajectory was vertically mirrored (the same trajectory could start moving to the left or to the right) on half of the repeats to minimize recognition or memorization.

Trajectories were randomly generated sum-of-sinusoid patterns generated from six individual sinusoidal components. Component frequencies always included the 0.05 Hz fundamental frequency, and then 5 randomly-sampled harmonic frequencies up to 0.4 Hz. Component amplitudes were sampled from the uniform distribution U(−1,1), and assigned in the order such that the largest magnitude amplitude went to the lowest frequency component and so forth until the highest frequency component was assigned the smallest magnitude amplitude. Trajectories were then scaled to a maximum horizontal deviation randomly sampled from the normal distribution N(12,1^2^) in degrees of visual angle. The variation in the maximum horizontal deviation was to introduce spatiotemporal uncertainty in the reversal points in the target trajectory. All components had a phase of 0 to ensure the trajectory started at the screen center. There were two additional criteria a trajectory had to satisfy: 1) the maximum speed had to be less than 30 deg/s, and 2) more than 75% of the time, the target had to be moving faster than 2 deg/s. These constraints were imposed to ensure the trajectory was likely to elicit smooth pursuit eye movements. If a sample trajectory did not meet these two criteria, it was discarded and resampled. This procedure resulted in fast, unpredictable, and varied trajectories that remained within the confines of the computer screen (see Figure 2 for an example trajectory).

**Figure 2:**
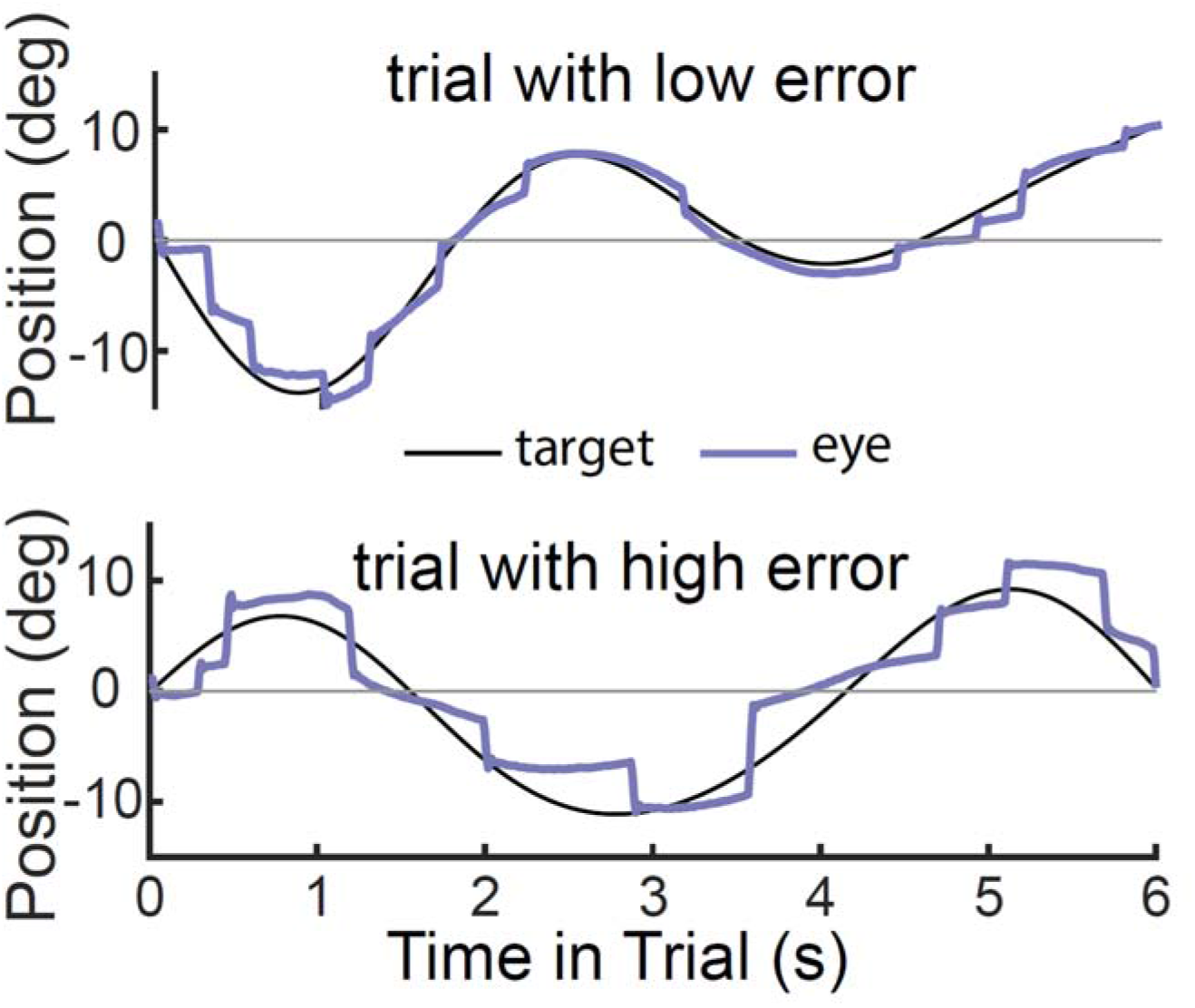
Example tracking behavior. The target trajectory was a sum-of-sinusoids (black smooth curve) and was shown to participants as a dot-cloud whose distribution was centered on the trajectory. The participant’s gaze (blue jagged curve) closely followed the target trajectory. Eye movements were a combination of periods of smooth tracking interspersed with saccadic jumps in position. **Top** shows an example trial with low tracking error, **Bottom** shows an example trial with higher tracking error chosen from one participant.

### Task

On each trial, participants performed two tasks. First, they were asked to track with their gaze the mean of four dots that were sampled from a Gaussian distribution centered on one of the 10 trajectories. The trajectory lasted 6 seconds and the dots were regenerated every frame at 60Hz. Then, participants were asked to judge whether their gaze tracking was better or worse than the average of all trials they performed so far. Tracking performance was defined to participants to reflect the spatial distance between the mean of the dots and their gaze at each time point of the trajectory.

The experiment was performed in two one-hour sessions held on different days. Each session began with a calibration procedure for the eye tracker, followed by a brief training task, and then the main task. The purpose of the training was to familiarize the participant with the stimuli and allow them to develop a sense of how well they can track the target. Participants performed 5 practice trials with no need to report their sensorimotor confidence afterwards. Participants were offered to repeat the training phase as many times as they wished before moving on to the main experiment. The trajectory set used during training was not the same as in the main task. In the main task, participants tracked the stimulus and then reported their confidence. In each session, participants performed 20 repeats of the 10 unique trajectories, half of which were horizontally mirrored, in a pseudorandom order. Thus, there were 40 repeats per trajectory total and 400 test trials in the whole experiment.

The trial structure was the same for the training and main tasks (see Figure 1). It began with a drift-correction, where the participant had to look at the screen center and press the space key before the trial would start. A blank screen was presented for 255 ms, after which the stimulus would appear at the screen center and move horizontally for 6 seconds. Then a blank screen was displayed and participants reported their confidence in their tracking performance as either “better” or “worse” than their average by pressing the right or left arrow key respectively. On the first trial, participants were given an onscreen reminder of the two confidence-response options. Small breaks were given every 10 trials in the main task for the participant to rest their eyes.

### Preregistration, data, and code availability

The code for running the experiment and performing the data analysis is available on Github at https://github.com/ShannonLocke/RepSinusoidTask, with repository release version 2 used for the data analysis reported within this manuscript. All raw and summary data for this experiment and its pilot version are available on the OSF repository at https://osf.io/j9asn/. Also included in this repository is the information about the target trajectories, additional results figures, pilot study report, and the data and code to generate the figures presented in this manuscript. This study was also pre-registered on OSF the 18th February 2022, with the full details available at https://osf.io/6s5gx.

### Hypotheses and Data Analyses

We preregistered four hypotheses and the matching analyses plan:

#### H1 (main)

Participants have above chance metacognitive sensitivity for sorting objectively better eye-tracking performance from objectively worse performance across the entire set of repeated trajectories (i.e., the AUROC is significantly greater than 0.5; see below and Locke, Mamassian, & Landy, 2020).

#### H2 (main)

Averaged across multiple repetitions of each trajectory, participants report “better” performance significantly more often for the 50% of repeats with lower tracking error than the 50% with higher tracking error for an individual repeated trajectory (i.e., a median split on objective error).

#### H3 (secondary)

The group-averaged temporal metacognitive sensitivity curve shows a recency effect (i.e., greater metacognitive sensitivity for later time bins).

#### H4 (secondary)

Metacognitive sensitivity for eye tracking is stable across days. Specifically, metacognitive sensitivity does not significantly differ between Session 1 and Session 2.

Pilot experiments revealed that participants performed the tracking task with a mixture of smooth pursuit and saccadic eye movements (see Figure 2 for an example tracking trace). We quantified tracking accuracy as the two-dimensional root-mean-squared-error (RMSE) between the gaze position and the tracking target. Both horizontal and vertical error contributed to tracking accuracy. To quantify the relationship between behavior and metacognition, we followed the approach developed by Locke, Mamassian, & Landy (2020). This involved collecting the distribution of tracking-error separately for trials labelled by the participant with a “better” confidence response and the trials labelled “worse”, and measuring the separation between these two distributions. A quantile-quantile comparison gives a receiver operating characteristic (ROC)-like curve, and the degree of separation in the confidence-conditioned distributions is reflected by the area under this curve (AUROC). An area of 0.5 indicates no metacognitive sensitivity, an area of 1 indicates perfect metacognitive sensitivity, and intermediate values indicate intermediate levels of metacognitive sensitivity. We used this approach to evaluate Hypotheses 1, 3, and 4. For Hypothesis 2, we used a novel approach to quantify metacognitive sensitivity that leverages the repeated-trajectories design of the current experiment. For each unique trajectory, we made a median-split of the repeats according to objective tracking accuracy: the half with better accuracy versus the half with worse accuracy according to the RMSE in the trial. Note for this analysis, we included all trials including those with RMSE scores that differed greatly from the participant’s mean accuracy. It is then possible to test if significantly higher confidence is given to the better half of repeats, for each trajectory.

### Exclusion criteria

Following our pre-registered protocol, we had to apply some exclusion criteria. We recruited 30 participants to reach our intended sample size of 27 participants. Three additional participants were recruited to replace two participants that only ever reported “better-than-average” tracking and a third participant who had a technical error occur in eye-tracking during recording, which we were able to correct later in post-processing. We detected an extreme confidence bias in 17 of the total 30 participants (56.7%), where the criteria was more than 75% of confidence responses favoring one the two choice alternatives. Thus, we report the results for the full sample (*n=30*) as well as the no-extreme bias sample (*n=13*) for all of the tested hypotheses. We also removed trials with problematic eye recordings using the criteria that the trial RMSE was +/-3 SD from that individual participant’s mean. This occurred rarely, with an average of 98.6% of trials kept for analysis per participant.

## Results

Participants tracked an unpredictable moving trajectory with a mixture of smooth pursuit and saccadic eye movements (see Figure 2 for an example tracking trace). Along our four preregistered hypotheses, we first look at the relationship between the reported performance and actual performance across all trials for an estimate of overall sensorimotor confidence. Then we will look at whether metacognitive sensitivity is influenced by the trajectory characteristics, by computing confidence within repetitions of the same trajectories. Finally, we will look how metacognitive sensitivity changed within a trial or across sessions.

### Overall metacognitive sensitivity (Hypothesis 1)

We first examined if participants had above-chance levels of metacognitive sensitivity for their eye-tracking behavior. We compared the distribution tracking accuracy for trials with reported better or worse performance and used that to compute an ROC-like curve (see Methods for more details). An area of 0.5 under this curve indicates no metacognitive sensitivity, an area of 1 indicates perfect metacognitive sensitivity, and intermediate values indicate intermediate levels of metacognitive sensitivity. In the full sample, we could compute the ROC-like curves for 28 of the 30 participants (see Figure 3), with the average area under the ROC-curve (AUROC) being 0.61 +/-0.02 SEM. The two remaining participants only ever responded “better-than-average” and so there was no “worse” error distribution available for the calculation. The level of metacognitive sensitivity in the full sample minus these two participants was significantly above chance (t(27)=6.69, p<0.05, Cohen’s d=1.27). This result remains unchanged if we exclude all participants with an extreme confidence bias (0.61 +/-0.02, t(12)=4.34, p<0.05, Cohen’s d=1.20; see Methods for more details).

**Figure 3:**
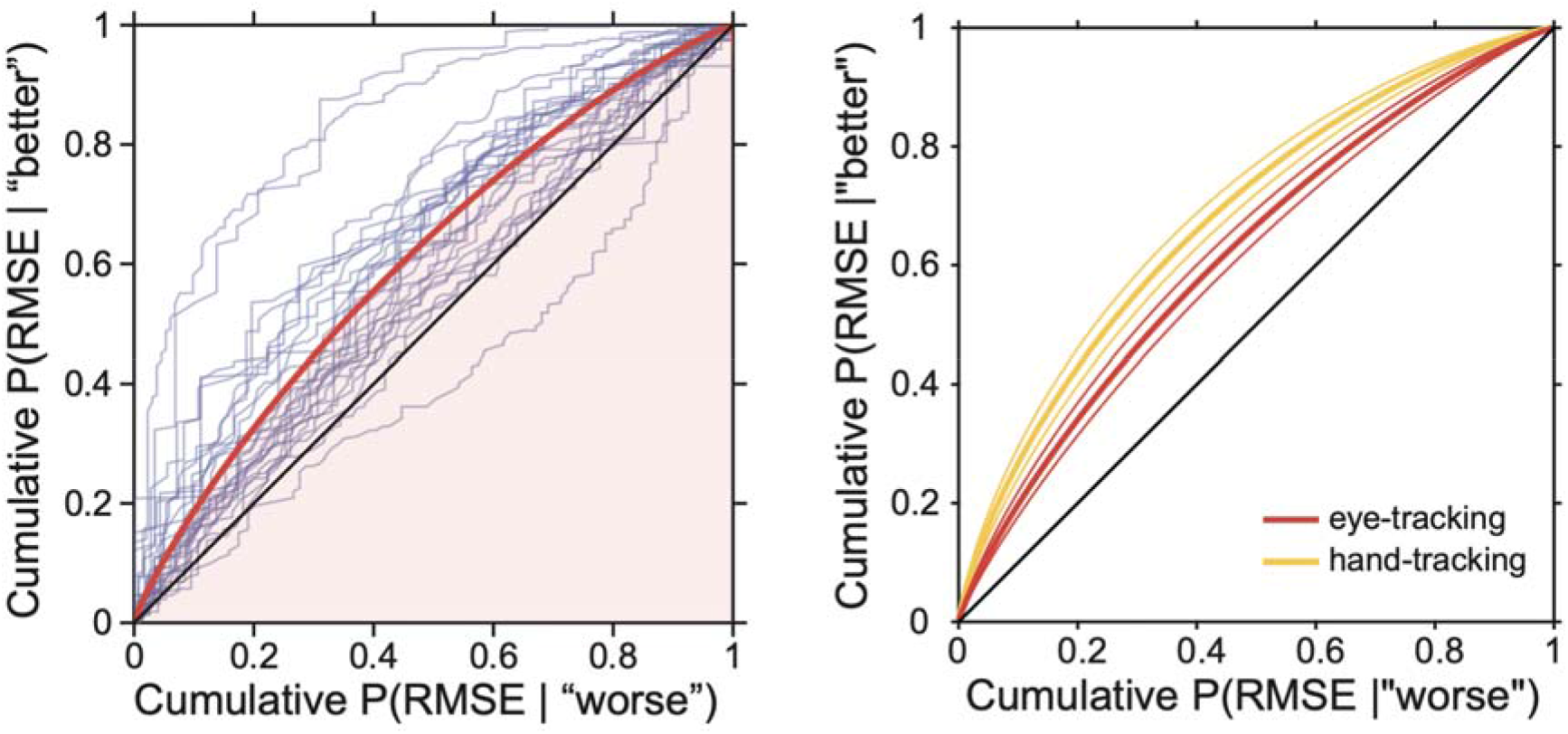
Metacognitive sensitivity results. **Left** An ROC-like analysis was applied to the confidence-conditioned tracking-error distributions for each participant (blue), with metacognitive sensitivity reflected by the area under the curve. Two participants had too few “worse” responses for this analysis and therefore are not shown. Tracking error was calculated as the root-mean-squared-error (RMSE) between gaze and target across the entire tracking period. Solid diagonal line: theoretical curve for a no-sensitivity observer. Red shading shows average metacognitive sensitivity across participants. **Right** red curve demonstrating the equivalent d’ for the current eye tracking experiment. The superior metacognitive sensitivity of the average observer in the hand-tracking study of Locke, Mamassian, & Landy (2020) is shown in yellow for comparison. Thin lines represent standard error the mean.

To compare the present results with the metacognitive sensitivity found for hand movements found in a previous study (Locke et al., 2020), we show the metacognition of the average participant in the present study using an equivalent d’ calculation (red curve in Figure 3) and contrast it against the previous results in visuomotor tracking (yellow curve). More precisely, the equivalent d’ is computed as follows: for a given empirical AUROC computed from two conditional error-distributions, that do not follow any particular standard probability distribution, the equivalent d’ is the separation in the means of two standard normal distributions that would result in the exact same AUROC value. Mathematically, the equivalent d’ is computed as follows: −−. The equivalent d’ for the present eye-tracking study is 0.39, which is much lower than the equivalent d’ of 0.66 found in the hand-tracking study of Locke, Mamassian, & Landy (2020).

### Repeated-trials analysis (Hypothesis 2)

A difficulty with the metacognitive-sensitivity analysis presented above is that it is unclear if there are some difficulty cues observable in each stimulus that participants can use to infer their tracking performance. For example, stimuli that move faster are often harder to track. This cue-based method of inferring tracking performance is in contrast to a more monitoring-based approach where the participant is genuinely keeping track of the errors they make as the trial progresses. One way we propose to eliminate potentially salient difficulty cues from the stimulus is to present the same stimulus multiple times. Tracking error will vary between these repeats, and we can measure if the confidence responses of the participant detect these performance variations (see Methods for more details). On average, participants were 8.21% +/-1.79% more likely to rate a more-accurate tracking trial as “better-than-average” in their confidence report. This pattern of higher confidence for more accurately-tracked trials was observable for all 10 unique trajectories (see Figure 4). This difference in confidence reports is significant (t(29)=4.57, p<0.05, Cohen’s d=0.83), and the results are unchanged if the extremely-biased participants are excluded from the analysis (13.27% +/-3.30%, t(12)=4.03, p<0.05, Cohen’s d=1.12). This supports a monitoring-based explanation of sensorimotor confidence for eye tracking.

**Figure 4:**
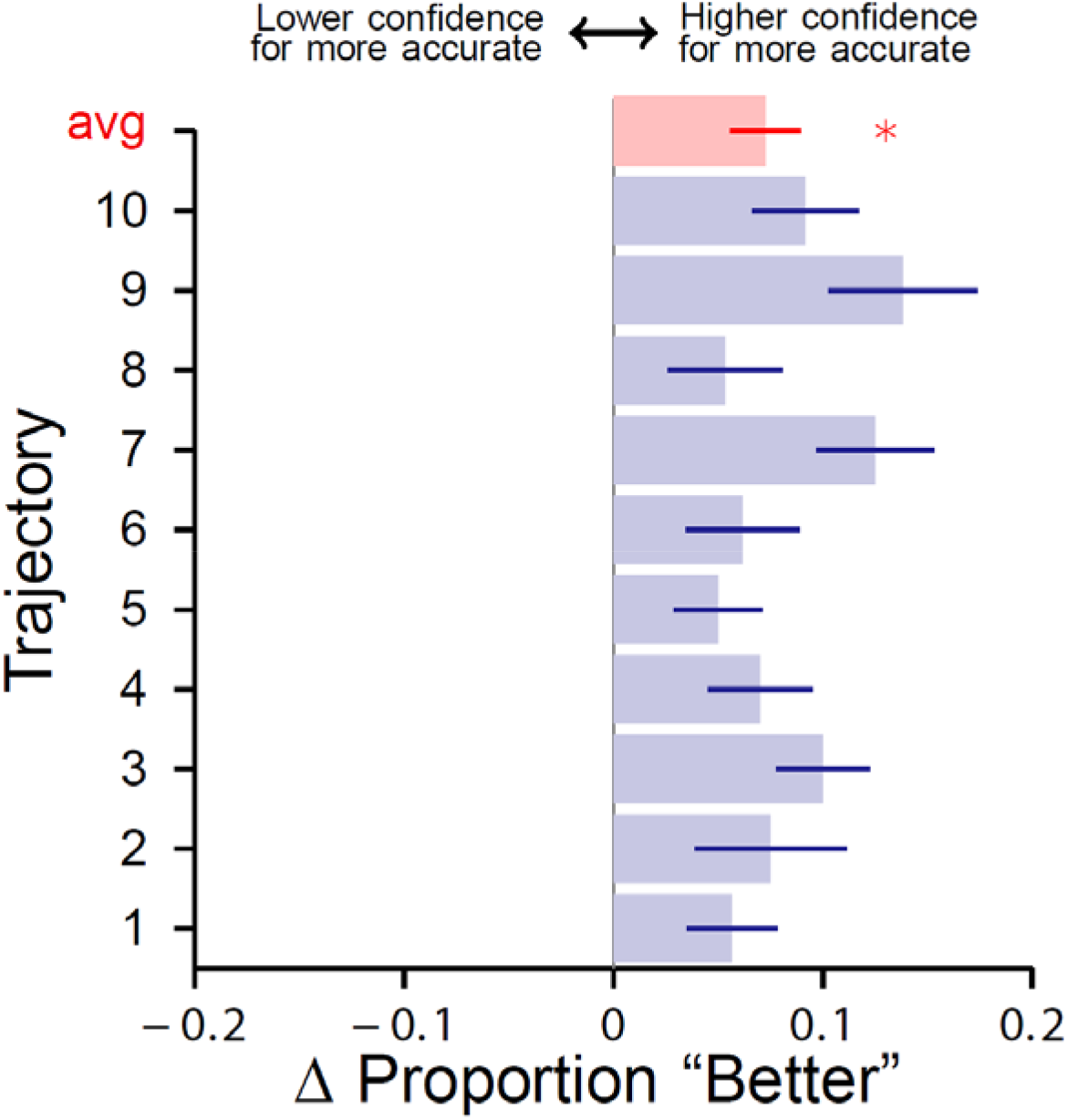
Results of the repeated-trials analysis. For all repeated trajectories, participants on average responded “better” more often for the half of trajectory repeats with the better tracking (i.e., lower RMSE). Individual trajectory results are shown in blue, and the average of all trajectories in red. Error bars: SEM.

### Temporal dynamics of metacognition (Hypothesis 3)

We also performed a temporal analysis of the metacognitive sensitivity of observers to test if there was a recency bias in metacognition. Previously, it has been shown that confidence in hand-tracked targets disproportionally relies on the tracking error at the end of the trial (Locke, Mamassian, & Landy, 2020). We followed the procedure of this previous study to compute the temporal metacognitive sensitivity curves by separately computing the AUROC using the error in each of the six 1-second time bins (see Figure 5A). We then tested for a recency effect by measuring if the average AUROC in the 5- and 6-second bins was significantly greater than the average AUROC of the 3- and 4-second bins. The first two seconds of tracking were ignored in this analysis as tracking error can take a couple of seconds to stabilize and we did not observe a systematic difference in tracking accuracy across the remaining trial. The AUROC was significantly higher by 0.05 +/-0.01 for the final two seconds as compared to the middle two seconds (t(27)=4.00, p<0.05, Cohen’s d=0.76) in the full population minus the two participants for which we could not compute their AUROC. However, this analysis did not reach significance in the sample excluding extremely-biased participants (0.03 +/-0.02, t(12)=1.60, p=0.13, Cohen’s d=0.44).

**Figure 5:**
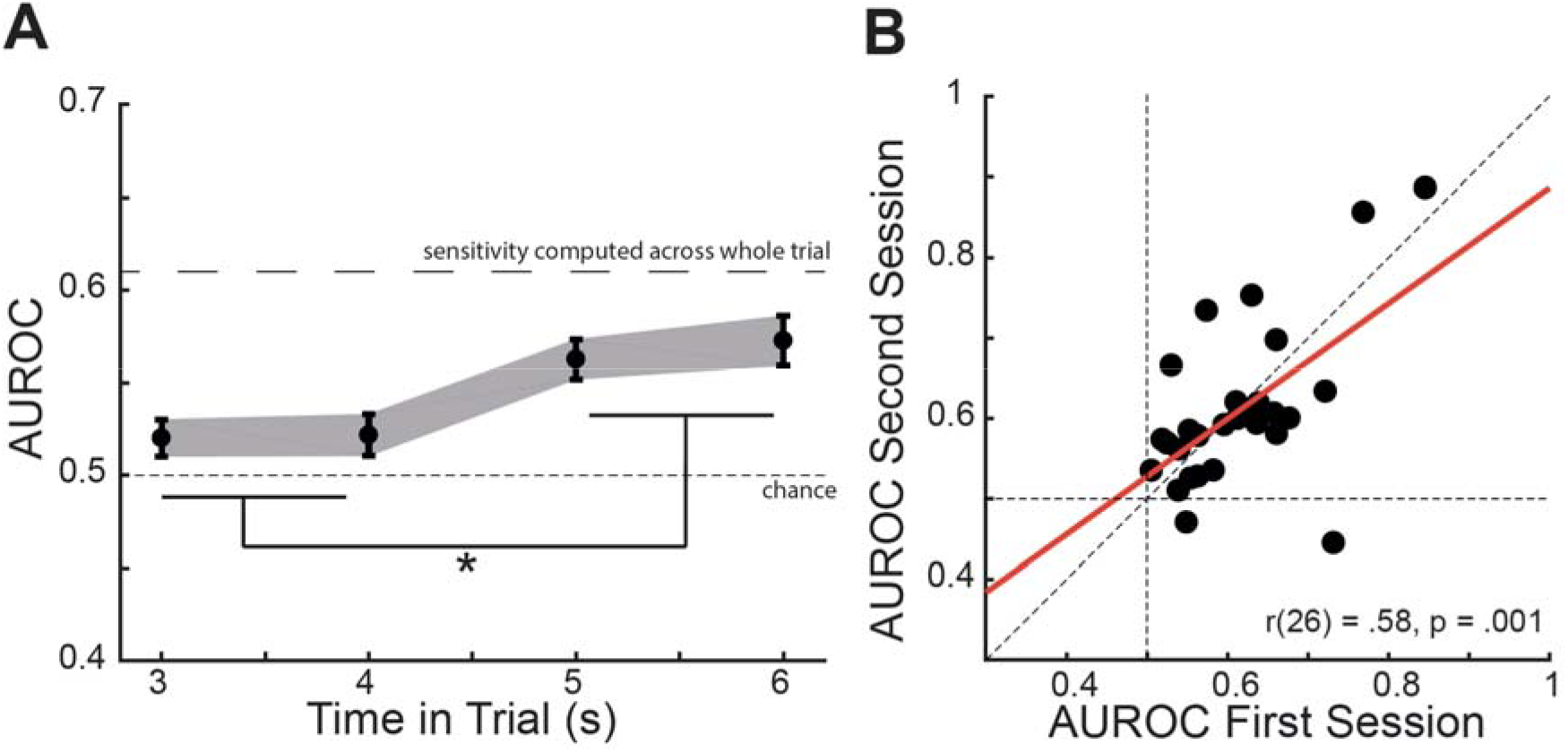
Temporal dynamics and stability of metacognitive sensitivity. **A** The average AUROC for all observers across time within in a trial. The AUROC is computed separately for each second of the trial while ignoring the first two seconds. The shaded area shows the SEM. **B** The AUROC for each participant, represented by separate dots, for the first and second sessions, which were separated by multiple days. Red line shows a linear regression fitted to the data.

### Metacognitive sensitivity across sessions (Hypothesis 4)

Our final analysis considered whether any metacognitive learning occurred between the first and second sessions of the experiment that were conducted on different days. To test this hypothesis, we computed the metacognitive sensitivity separately for each session. For both the larger sample and the extreme-bias-excluded sample, we did not find any significant difference in metacognitive sensitivity between the sessions. The differences were -0.004 +/-0.02 (t(27)=-0.27, p=0.79, Cohen’s d=-0.05) and -0.01 +/-0.01 (t(12)=-0.76, p=0.46, Cohen’s d=-0.21) respectively. This indicates that metacognitive sensitivity was stable across the experiment. As an additional exploratory measurement, instead of focusing on the average, we also looked at the individual variation in metacognitive sensitivity (see Figure 5B). We observed that sensitivity stayed comparable (r(26) = .58, p =.001), suggesting temporally stable differences in sensitivity across participants.

## Discussion

The goal of the present study was to test whether humans are capable of actively monitoring the performance of their own eye movements. Across all trials we found a low, but significant agreement between the reported and actual tracking performance of observers, demonstrating sensorimotor confidence for eye movements (Figure 3). When we calculated the metacognitive sensitivity separately for the multiple repeats of each of the different target trajectories (Figure 4), sensitivity was present for each of the trajectories suggesting that the effect was not related to stimulus characteristics. When looking into the temporal evolution of the effect, we observed that the reports of performance were mostly related to tracking performance late in the trial (Figure 5A), but that overall sensitivity was comparable and stable across different sessions measured on separate days (Figure 5B).

### Heuristics or performance monitoring

While our results suggest a limited, but successful performance monitoring, the information was used inefficiently. One factor that needs to be carefully considered is how other potential and even irrelevant cues could have affected the subjective reports. It is known that confidence reports can be affected by heuristics and assumptions about the task (Mole et al., 2018; Navajas et al., 2017; Spence et al., 2016). For example, despite similar performance, participants are biased to report higher confidence if the task involves their own active movement in comparison to a passive movement or a visual task (Charles et al., 2020). Confidence can even be manipulated independently of performance, for example when motion characteristics (de Gardelle & Mamassian, 2015) or stimulus visibility (Maniscalco et al., 2016; Rausch et al., 2015) are changed across trials. For our current results, we believe it is unlikely that assumptions about the task or differences in target visibility can explain our results, since also our AUROC is not affected by potential biases (similarly to d’ in signal detection theory). While there were different target trajectories, we found consistent and stable sensitivity for each of them when directly comparing the trials with good and bad tracking performance. If observers would have used heuristics about the difficulty of individual trajectories to make their confidence judgements, we would expect mainly positive reports for some of the trajectories and mainly negative reports for other trajectories. However, our repeated-trajectories analysis does not support this, and instead suggests that the reports were based on active monitoring of tracking performance. Furthermore, if observers would have based their decisions on simple heuristics or sensory information about the target trajectory, there should not have been a stronger influence of tracking error towards the end of the trial. This recency effect demonstrates that the subjective report was related to performance monitoring and that some moments were treated differently than others (Locke et al., 2020). Locke and colleagues could show that such a recency effect also occurs for unpredictable trial durations and interpret it as a signature of the temporal accumulation of a confidence signal with “leaky accumulation” and memory-limitations (see their paper for a more detailed discussion). Together, these points suggest, while heuristics can play an important role for confidence judgements, in our task metacognitive sensitivity seems to be mainly driven by an inefficient, but successful performance monitoring.

### Confidence across sensory and sensorimotor processing

So far, previous studies mainly have considered the contribution of movements in perceptual confidence (Fleming & Daw, 2017; Kiani et al., 2014; Yeung & Summerfield, 2012), or used motor behavior as index of perceptual confidence (Dotan et al., 2018; Patel et al., 2012; Resulaj et al., 2009). However, these studies mostly focused on simple behavior and did not include an assessment of the potential role of additional motor variability. Extending the work from Locke and colleagues (2020) on hand movements, we show that people also can make successful confidence judgements about eye movements and their variability. This supports the idea that confidence could serve as a common currency to evaluate performance across tasks (Ais et al., 2016; de Gardelle et al., 2016; Faivre et al., 2018). Confidence judgements are correlated when judging performance for a task that involves judging the position of an active movement and a simple perceptual passive viewing task (Charles et al., 2020). In line with measures of perceptual confidence (Ais et al., 2016; de Gardelle & Mamassian, 2015; Navajas et al., 2017), we also observed stable individual differences in the sensitivity of observers that stayed consistent across multiple days. Thus, it would be interesting to test whether sensorimotor confidence is also linked to other instances of confidence measurements.

### Differences in sensitivity between eye and hand movements

When comparing metacognitive sensitivity measured for eye movements with the previous work using a similar paradigm for hand movements (Locke et al., 2020), sensitivity for eye movements was lower (Figure 3). When comparing the two paradigms directly, there was a potentially important difference: In the hand movement task, there was an additional visual representation of the hand position on the screen, which could have allowed for a more direct assessment of tracking performance. We choose to not have a similar representation based on the gaze position since the target would have otherwise been an additional salient and dynamic distractor that would have been constantly aligned with the gaze position and therefore would have interfered with tracking performance. It was proposed that motor awareness is restricted to only the initiation of actions and evaluation of outcomes, with only limited possibilities to monitor the movement itself (Blakemore et al., 2002; Blakemore & Frith, 2003; Haggard, 2017). It is difficult to assess where the initiation and evaluation of a movement occur for a continuous tracking target as the one used in our experiment. However, having the additional target representation and a comparison to the current gaze position, the evaluation of the hand movement should have been easier, which could explain the higher sensitivity.

Next to the difference in performance feedback, an additional explanation could be that there might be a difference in the quality of evaluating and monitoring eye and hand movements. While eye (Spering & Carrasco, 2015; Tavassoli & Ringach, 2010) and hand movements (Fourneret & Jeannerod, 1998; Goodale et al., 1986; Prablanc & Martin, 1992) can be controlled by information we are perceptually unaware of, there seems to be a particular unawareness about how we move our eyes (Goettker et al., 2018; Nieuwenhuis et al., 2001; van Zoest & Donk, 2010). Such a difference could be based on the differences in control: While eye movements are controlled by only a few muscles and do not need much energy, successful hand movements require much more energy and force. It is also the case that while eye movements only increase our understanding of the environment, hand movements can modify it; and sometimes produce injuries from contact with the environment. Therefore, there might have been greater evolutionary pressure to be better at monitoring one’s hand than eye movements.

Additionally, while we observe metacognitive sensitivity for the eye, it might not be based on a judgement of actual motor performance, but a judgment of a more general state. Earlier work showed that humans are able to estimate their own saccade latency reasonably well (Vencato & Madelain, 2020), however such a performance could have been achieved without actually knowing about when the eye movement began. Instead, participants could be judging the state of attention and preparedness at the beginning of the trial (Tomassini et al., 2015; VanRullen, 2016). Similarly, knowing the current level of attention could serve as a proxy for the quality of the movement, without actually having access to the movement signal. In an extreme case, one could achieve a good metacognitive sensitivity by just purposefully showing bad performance in half of the trials and judging those as bad and the others as good. Since our estimated sensitivity for eye movements was rather low, participants clearly did not follow this strategy, but potentially a more implicit, unconscious version of the same idea was present. The recency effect could also be explained by simply focusing on the level of concentration people were paying towards the end of the trial. The difference between the sensitivity for eye and hand movements could then be based on the fact, while for both tasks there is an overall sensitivity for current state of the observer, for hand movements additional monitoring and evaluation processes were available.

## Conclusion

Our goal was to understand whether observers can judge the accuracy of their own eye movements. We found across all trials and even within repetitions of the same trajectory, participants did show a sensitivity for distinguishing good from bad trials, thus displaying metacognitive sensitivity. The observed sensitivity was mostly related to performance towards the end of the trial, which matched previous reports when judging the quality of hand movements, although overall sensitivity for eye movements was significantly lower. These results provide an additional piece of evidence for sensorimotor confidence, and open interesting questions about why it differs across movements and how it could be related to other instances of confidence.

## Acknowledgments

The authors would like to thank Nina Küpke for her help in data collection. S.L. & P.M. were supported by an Anneliese Maier Award from the Alexander von Humboldt Foundation awarded to PM. A.G. & K.G. were supported by the Deutsche Forschungsgemeinschaft (DFG; project number 222641018-SFB/TRR 135 Project A1).

## Author contributions

Conceptualization, S.L., A.G., K.G. & P.M.; methodology, S.L. and A.G.; data collection, S.L. & A.G.; formal analysis, S.L.; writing—original draft, A.G. and S.L.; writing—review & editing, A.G., S.L., K.G. & P.M.; visualization, S.L. and A.G.; funding acquisition, K.G. and P.M.

